# rawR - Direct access to raw mass spectrometry data in R

**DOI:** 10.1101/2020.10.30.362533

**Authors:** Tobias Kockmann, Christian Panse

## Abstract

The Bioconductor project has shown that the `R` statistical environment is a highly valuable tool for genomics data analysis^1^, but with respect to proteomics we are still missing low level infrastructure to enable performant and robust analysis workflows in `R`. Fundamentally important are libraries that provide raw data access. Our `R` package `rawDiag` has provided the proof-of-principle how access to mass spectromerty raw files can be realized by wrapping vendor-provided APIs, but rather focused on meta data analysis and visualization^2^. Our novel package `rawR` now provides complete, OS independent access to all spectral data logged in Thermo Fisher Scientific raw files. In this technical note we present implementation details and describe the main functionality provided by the `rawR` package. In addition, we report two use cases inspired by real-word research task that demonstrate the application of the package.

**Availability:** https://github.com/fgcz/rawR

## Introduction

Mass spectromerty-based proteomics and metabolomics are the preferred technology to study the protein and metabolite landscape of complex biological systems. The orbitrap mass analyzer is one of the key innovations that propelled the field by providing HRAM data on a chromatographic time scale. Driven by the need to analyze the resulting LC-MS data, several specialized software tools haven been developed in the last decade. In the academic environment, [MaxQuant](https://maxquant.org/) and [Skyline](https://skyline.ms/project/home/begin.view?) are beyond the most popular ones. These software tools usually offer GUIs that control running predefined analysis templates/workflows including free parameters that need to be defined by the user. In parallel, projects like [OpenMS](https://www.openms.de/) or [pytemocis](https://github.com/levitsky/pyteomics) developed, but chose a fundamentally different approach. They aim at providing software libraries bound to specific programming languages like `C++` or `Python`. These naturally offer greater analytical flexibility, but require programming skills form the end user and have therefore not reached the popularity of their GUI counterparts. Proteomics and metabolomics specific libraries have also been developed for the [R](https://www.r-project.org/) statistical environment, but these mainly support high-level statistical analysis, once the raw measurement data has undergone extensive pre-processing and aggregation by external software tools (often the GUI-based once listed above). A typical example is the `R` package [MSstats](http://msstats.org/) for the statistical analysis of LS-MS experiments with complex designs or [MSqRob](https://github.com/statOmics/MSqRob). MSstats can process MaxQuant or Skyline output and creates protein/peptide level estimates whether the biological system shows statistically significant regulation. In a nutshell, these tools provide statistical post processing. Libraries that support working with the spectral data in `R` also exist, for instance the BioC package [MSnbase](https://www.bioconductor.org/packages/release/bioc/html/MSnbase.html), but require conversion of raw data to exchange formats like [mzML](http://www.psidev.info/mzML). These conversion are primarily supported by the [ProteWizard](http://proteowizard.sourceforge.net/) project and its software tool `MSconvert`.

We strongly believe that a library providing raw data reading functionally would finally close the gap and facilitate modular end-to-end analysis pipeline development in `R`. This could be of special interest to research environments/projects dealing with either big data analytics, or to scientists that are interested in code prototyping without having a formal computer science education. Another key aspect regarding multi omics integration of proteomics and metabolomics data is the fact that high-throughput genomic data analysis is already done mostly in `R`. So proteomics and metabolomics could finally “join the party”! This is primarily due to the [BioC project](https://www.bioconductor.org/) that currently provides >1900 open source software packages, training & teaching, and a very active user & developer community^1^. Having these thoughts in mind we decide to implement our R package `rawR`. `rawR` utilizes a vendor-provided API to access spectral data logged in proprietary raw files. These binary files are written by all orbitrap mass spectrometers, unlocking an incredible amount of the global LC-MS data, also stored in public repositories like [ProteomeExchange](http://www.proteomexchange.org/). In this manuscript, we present a first package version/release and show case its usage for bottom-up proteomics data analysis.

## Implementation

Our implementation is build on two layers, the `R` and the `C#` layer which exchange information using file i/o. The following section describes the package topology from top-down, starting with a rational for the object design.

`R`

Mass spectrometry (MS) typically uses two basic data items: mass spectra and chromatograms.

### Spectra

All mass spectra are recorded by scanning detectors (mass analyzers) that log signal intensities for ranges of mass to charge ratios (m/z), also referred to as position. These recordings can be of continuous nature, so called profile data (p), or appear centroided (c) in case discrete information (tuples of position and intensity values) are sufficient. This heavily compacted data structure is often called a peak list. In addition to signal intensities, peak list can also cover additional peak attributes like peak resolution (R), charge (z), or local noise estimates.

### Chromatograms

Chromatograms come in different flavors, but are always signal intensity values as a function of time. Signal intensities can be point estimates from scanning detectors or plain intensities from non scanning detectors (e.g. UV trace). Scanning detector (mass analyzers) point estimates can be defined in different ways by for instance summing all signals of a given spectrum (total ion chromatogram or TIC), or by extracting signal around an expected value (extracted ion chromatogram = XIC), or by using the maximum signal contained in a spectrum (base peak chromatogram = BPC). On top, chromatograms can be computed from pre-filtered lists of scans. A total ion chromatogram (TIC) for instance is typically generated by iterating over all MS1-level scans.

We therefore decided to implemented corresponding objects following `R`s `S3` OOP system named `rawRspectrum` and `rawRchromatogram` that closely resemble the above definitions^3^. The package provides functions to create and validate class instances (objects), but typically instances are generated by reader functions that expect rawfiles as input data (see Table 1. for an overview). We refer to collections of objects as sets (e.g. `rawRspectrumSet`). The constructors (`rawRspectrum()` and `rawRchromatogram()`) primarily exist for (unit) testing proposes, or for simulating data (no binary input data is required). One can for instance generated spectra showing predefined patterns (ion series derived from a peptide sequence) or sample from base `R`s collection of distributions (in preparation). We also implemented basic generics for printing and plotting of objects that we use for visualization throughout this manuscript. To minimize dependencies, we choose to stick to `base R`, but it would be relatively easy to write your own plotting functions using `lattice` or `ggplot`. The reader functions typically call compiled `C#` code (see next section). We call these `C#` methods wrappers.

**Table 1:**
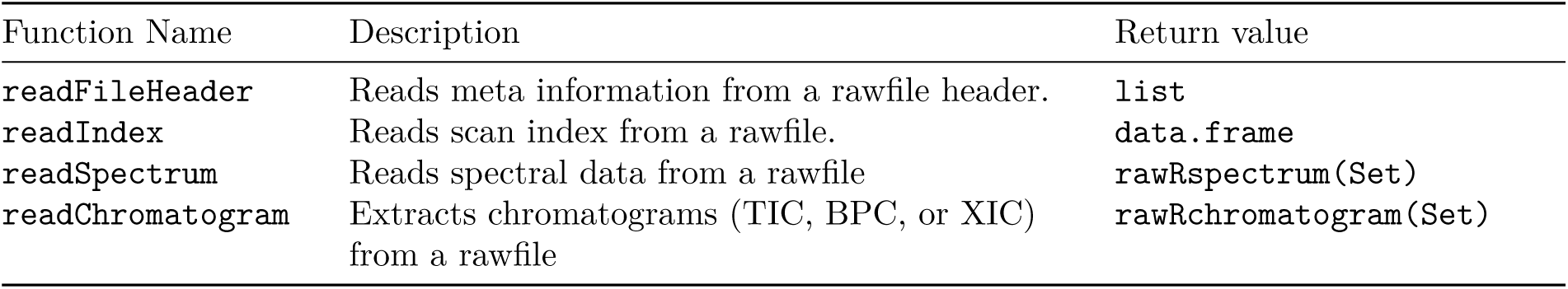
lists the most important `rawR` package functions connected to reading functionality. More details can be found in the package documentation.

`C#`

The `base::system2` R function invokes methods compiled in the compiled `rawR.exe` .NET application. `rawR.exe` utilizes the NewRawFileReader .Net assembly provided by Thermo Fisher Scientific^4^. Therefore, it can read the proprietary vendor files and pass R code for S3 class R^5^ objects back to the R layer of `rawR`. The `C#` source code and a precompiled binary is shipped with the released R package and runs on Linux, Microsoft Windows, and MacOSX on the X86_64 hardware architecture. On Linux and MacOS the [mono](https://www.mono-project.com/) compiler is required to run the wrapper functions.

### Example binary data

The [example file](http://fgcz-ms.uzh.ch/~cpanse/20181113_010_autoQC01.raw) (MD5: `a1f5df9627cf9e0d51ec1906776957ab`) used throughout this manuscript contains fourier-transformed orbi trap spectra (FTMS) recorded on a Thermo Fisher Scientific Q Exactive HF in positive mode (+). The mass spectrometer was operated in line with a nano UPLC and a nano electrospray source (NSI). MS2 spectra were generated by HCD fragmentation at a normalized collision energy (NCE) of 27. All spectra were written to disk after applying centroiding (c) and lock mass correction. The analyzed sample consists of the iRT peptide mix (Biognosys) in a tryptic BSA digest (NEB) and was separated applying a 20 min linear gradient on C18 RP material at a constant flow rate of 300 nl/min.

Additional raw data for demonstration and extended testing is available through the BioC data package [tartare](https://bioconductor.org/packages/release/data/experiment/html/tartare.html). Lions love raw meat!

## Results

The following sections are inspired by real-life research/infrastructure projects, but have been stripped down to the bare scientific essentials in order to put more weight on the software application. We display source code in grey shaded boxes including syntax highlights. Corresponding `R` command line output starts with `##` and is shown directly below the code fragment that triggered the output. All figures are generated using the generic plotting functions of the package.

### Use Case I - Analyzing orbi trap spectra

The orbitrap detector has been a tremendous success story in MS, since it offers high resolution, accurate mass (HR-AM) data on a time scale that is compatible with chromatographic analysis (LC-MS). It is therefore heavily used for bottom-up proteomics, but analyzing orbitrap data in `R` has so far only been possible after raw data transformation to exchange formats like mz(X)ML. This use case shows how easy it is to work directly with the binary raw data, after installing our `R` package `rawR` that applies vendor APIs for data access. For demonstration purposes we use a complete LC-MS run recorded on a Q Exactive HF Orbitrap. The 35 min run resulted in 2.1881 × 10^4^ scans that were written to disc. Already type setting the above lines uses `rawR` functionality, since the instrument model, the time range of data acquisition, and the number of scans is extracted from the binary file header (Note: This manuscript was written in `R markdown` and combines `R` code with narration). The respective function is called `readFileHeader()` and returns a simple `R` object of type `list` (see Table 1).

Individual scans or collection (sets) of scans can be read by the function `readSpectrum()` which returns a `rawRspectrum` object or `rawRspectrumSet`. Our package of course also provides generics for printing and plotting these objects. The following code chunk depicts how a set of scans is read from the rawfile and the corresponding Figure 1 shows the resulting plot for scan `9594`:

**Figure 1:**
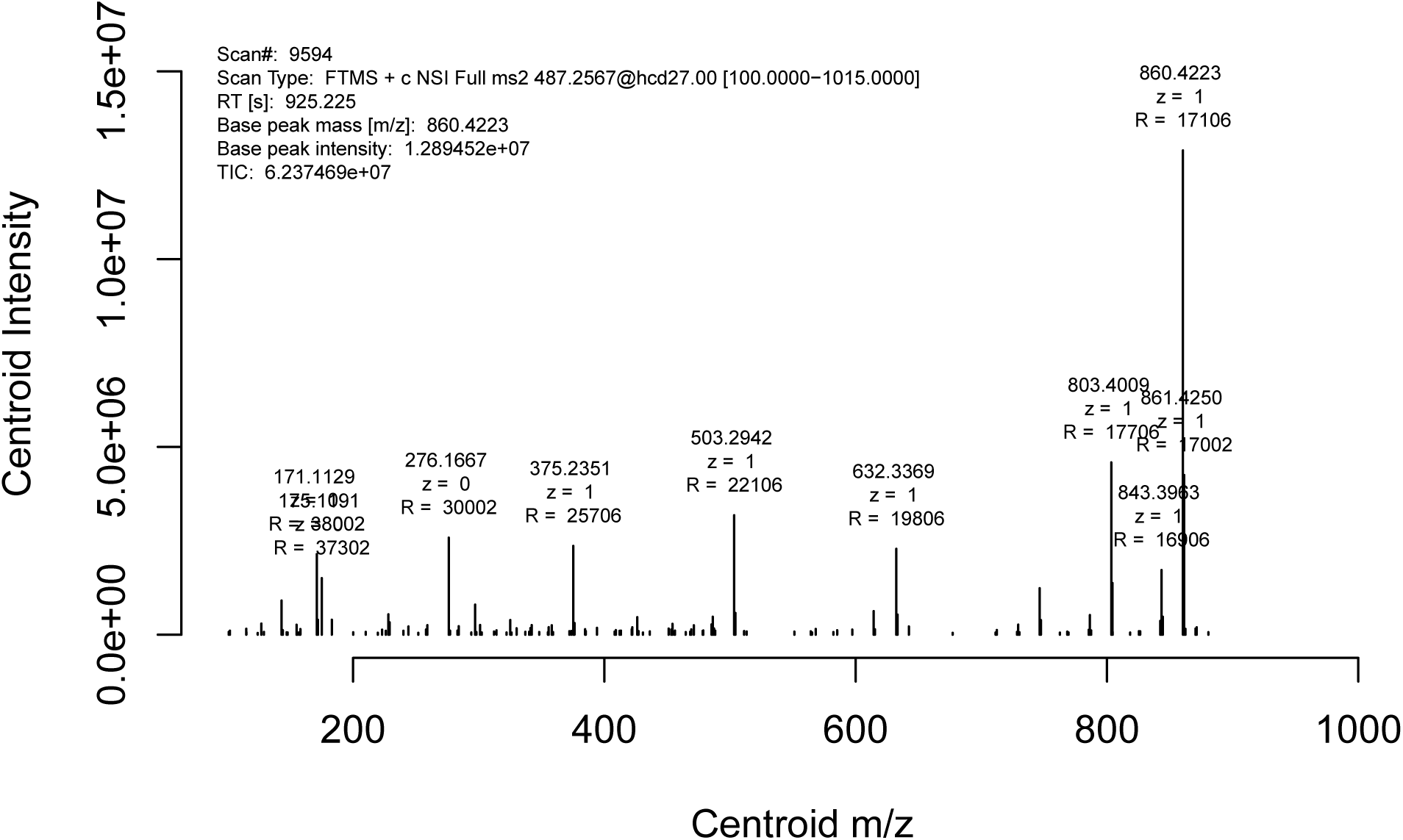
Plot of scan number 9594 showing a centroided tandem mass spectrum of the iRT peptide precursor LGGNEQVTR++ in positive mode. The scan was acquired on an orbitrap detector incl. lock mass correction and using a transient of 64 ms (equal to a resolving power of 30’000 at 200 m/z) and injection of 100’000 charges (AGC target). Peak attributes like m/z, charge (z), and resolution (R) are shown above the peaks.

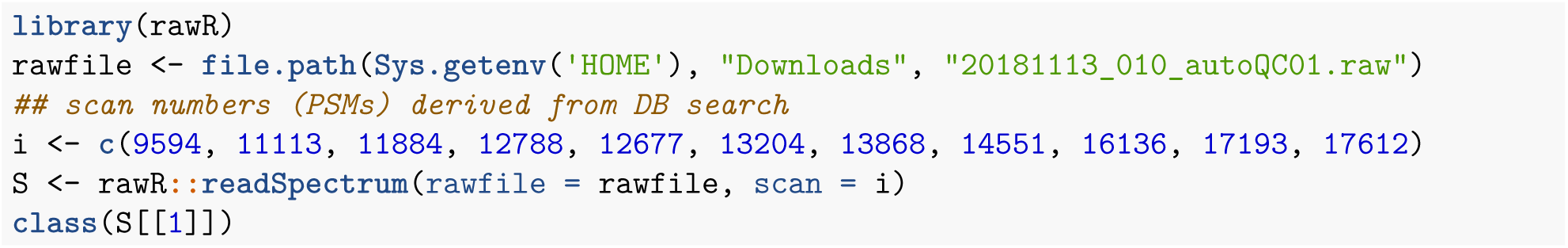

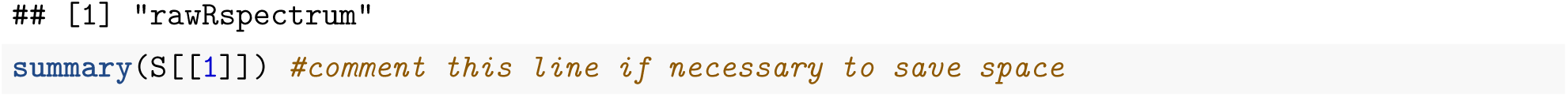

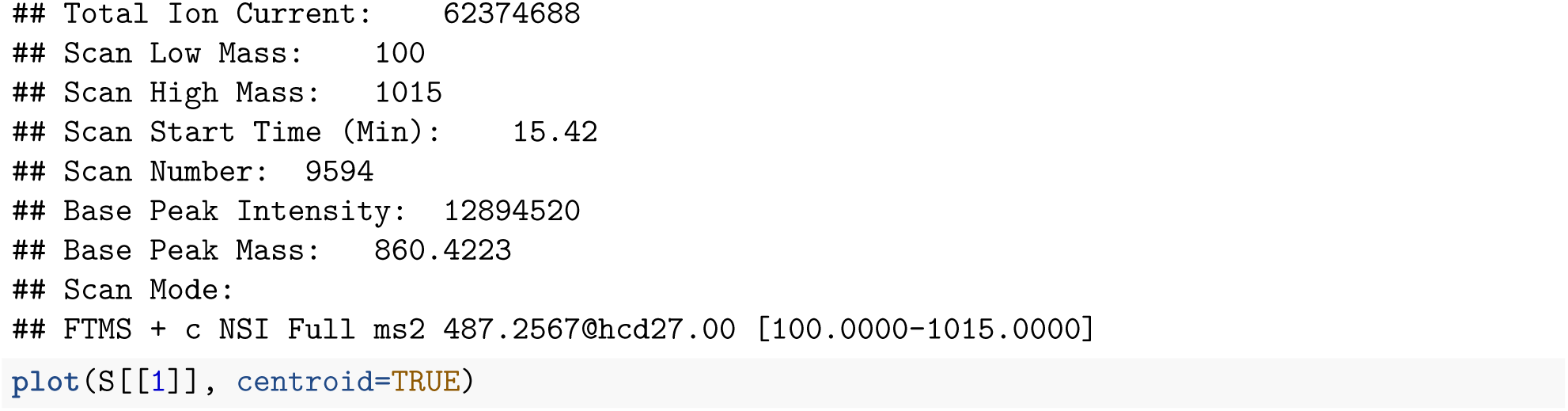

The plot shows typical orbitrap peak attributes like resolution (R) and charge (z) above the most intense peaks when centroided data is available and selected. Centroided data also makes it possible to graph spectra using signal-to-noise as response value. This is potentially interesting, since orbitrap detectors follow

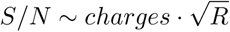

and signal-to-noise makes judging the signal quantity more intuitive than using arbitrary signal intensity units. Figure 2 shows that all fragment ion signals are several ten or even hundred fold above the local noise estimate.

**Figure 2:**
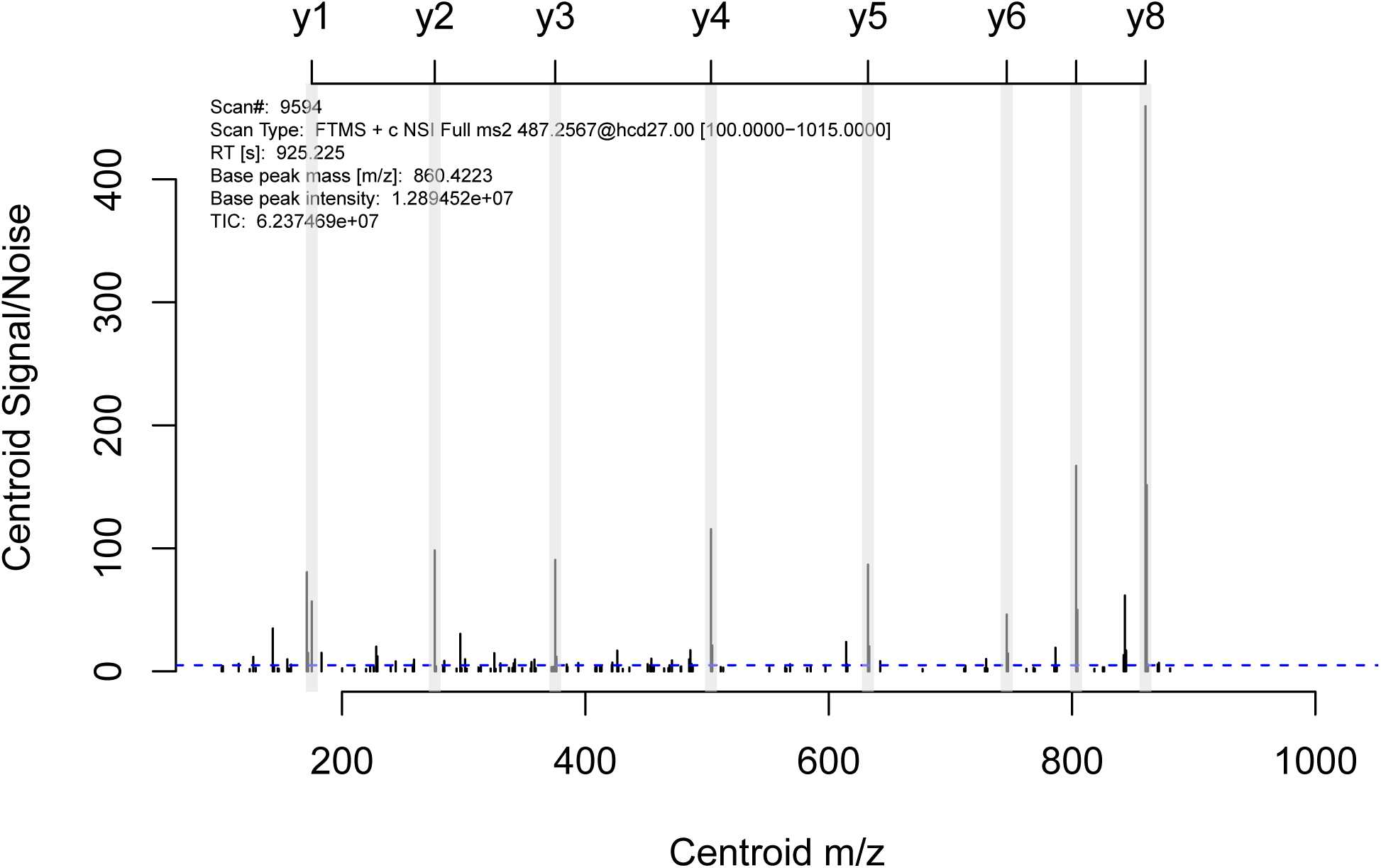
Spectrum plot using Signal/Noise option. The vertical grey lines indicate the *in-silico* computed y-ions of the peptide precusor LGGNEQVTR++ as calculated by the [protViz](https://cran.r-project.org/package=protViz) package^6^.

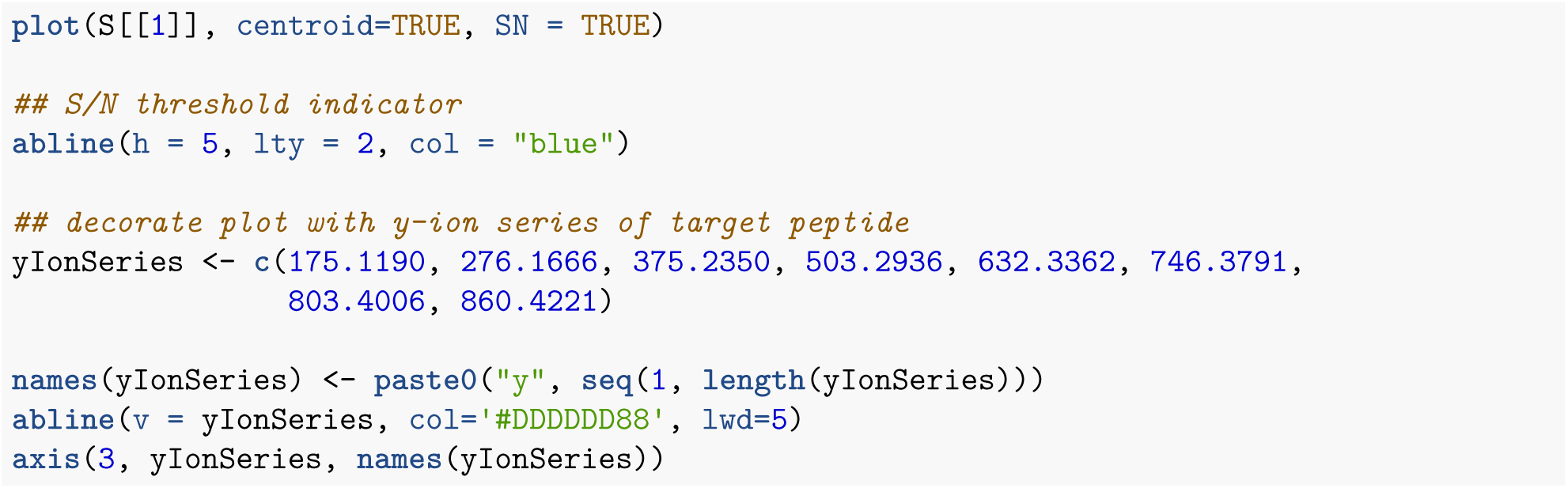

More sophisticated analysis workflows applying `rawR` functionality have been described in^7^. In short, marker ions found in HCD MS2 spectra for ADP-ribosylated peptides were annotated and cross compared at different collision energies. You may have notices that such things become relatively simple, since the `rawRspectrum` object provides easy access to normalized and absolute collision energies. A small molecule application using UVPD dissociation is described in^8^.

### Use Case II - iRT regression for system suitability monitoring

By applying linear regression one can convert observed peptide retention times (RTs) into dimensionless scores termed iRT values (iRTs) and *vice versa*^9^. This can be used for retention time calibration/prediction. In addition, fitted iRT regression models provide highly valuable information about LC-MS run performance. In this example we show how easy it is to perform iRT regression in `R` by just using the raw measurement data, our package `rawR`, and well known `base R` functions supporting linear modeling. To get a first impression of the data we calculate a TIC using the `readChromatogram()` function. Plotting the TIC shows chromatographic peaks between 15 and 28 min that could be of peptidic origin (Hint: There is also a `type = “bpc”` option if your prefer a BPC):

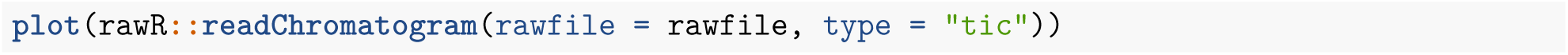

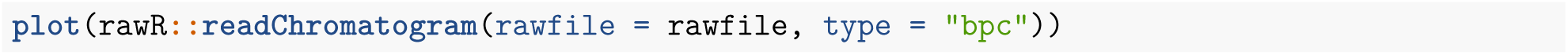

The initial step of iRT regression is now to estimate the empirical RTs of a peptide set with known iRT scores. In the simplest case, this is achieved by computing an extracted ion chromatogram (XIC) for iRT peptide precursors, given they were spiked into the sample matrix prior to data acquisition. Luckily ;-), our example data is iRT peptides in a tyrptic digest of BSA. The code chunk below demonstrates how the function `readChromatogram()` is called on the R command line to return a `rawRchromatogramSet` object of the type `xic`. This object is plotted for visual inspection.

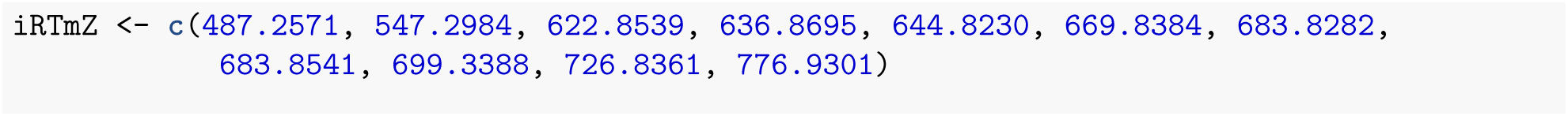

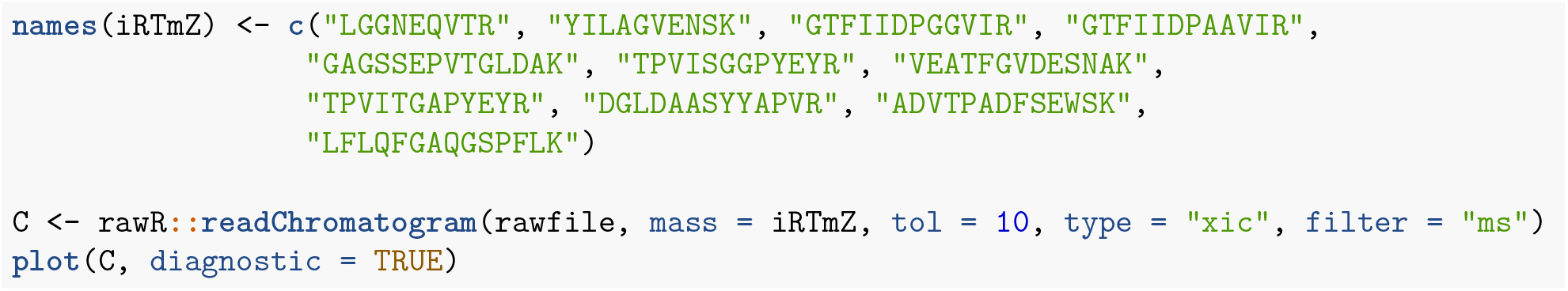

Be reminded that the intensity traces are not computed within `R` for instance by reading all scans of a raw file and subsequently iterating over a scan subset (This would be a greedy, but slow solution!). Instead, traces are directly calculated by a `C#` method (reference method code) that calls the vendor API. The API takes care of the filtering process (checks filter validity and applies the filter). On the `R` level there is no need to know *a priori* which scans match the filter rule, or implement vectorized operations (we generate multiple XICs simultaneously here). Only the API-returned output needs to be parsed into `rawRchromatogram` objects. By changing the filter, one can easily switch between generating precursor traces and fragment ion traces. The following code chunk shows how to create fragment ion chromatograms (y6 to y8) generated from scans that target LGGNEQVTR++:

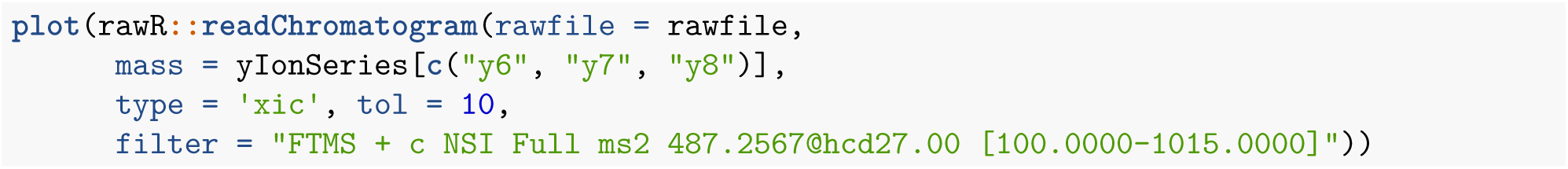

You will immediately recognize that this means our example data was actually recorded using parallel reaction monitoring (PRM), since 487.2567 was targeted in regular spaced intervals. You could confirm this by using the `readIndex()` function which returns a `data.frame` that indexes all scans found in a raw file and subsetting it for the scans of interest. The delta between consecutive scans is always 22 scans:

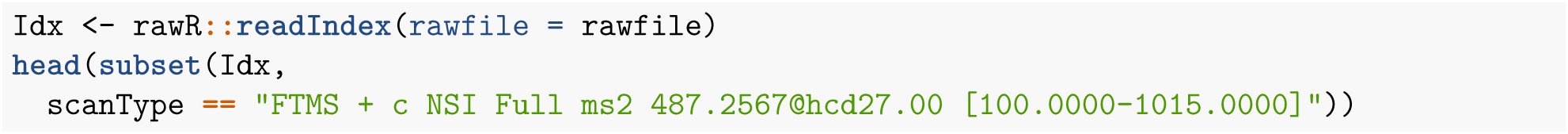

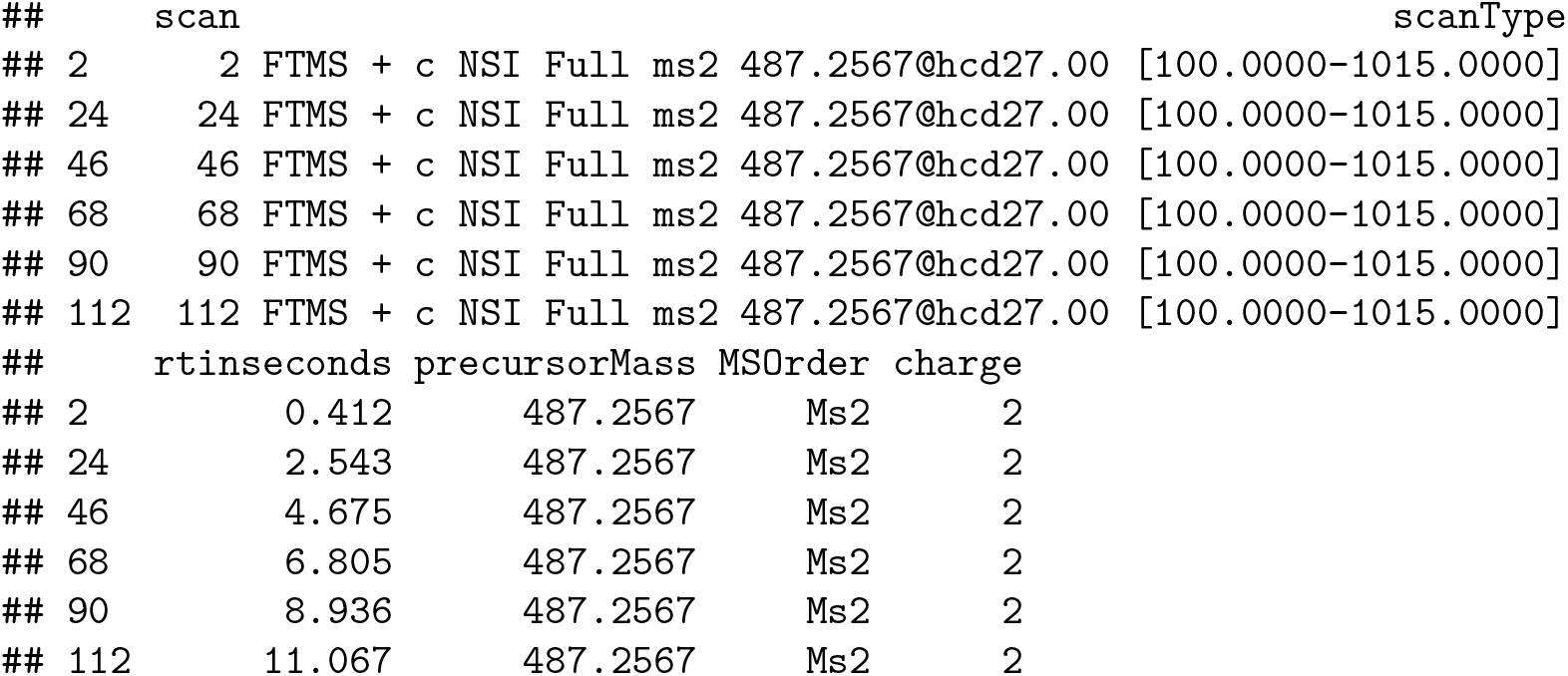

For regression, we now extract the RTs at the maximum of the intensity traces stored in the chromatogram object and fit a linear model of the form:

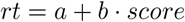

In theory, we could do this at the precursor or fragment ion level. For simplicity we show only the first option.

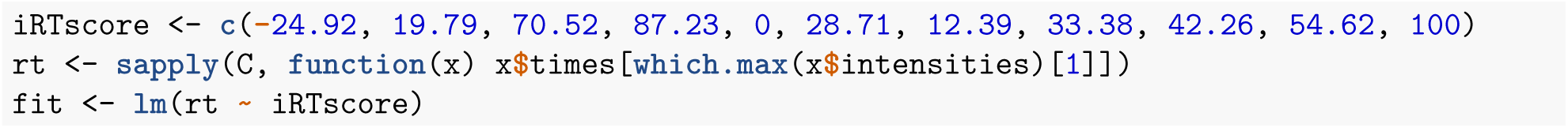

The fitted model can than be inspected using standard procedures. The output of code chunk `iRTscoreFitPlot`, in Figure 6, shows a visual inspection by plotting observed RTs as a function of iRT score together with the fitted model regression line. The corresponding R squared indicates that the RTs behave highly linear! This is expected since, the iRT peptides were separated on a 20 min linear gradient from 5 %B to 35 %B using C18 RP material (the change rate is therefore 1.5 %B / min). The magnitude of the slope parameter (b) is a direct equivalent of this gradient change rate. The intercept (a) is equal to the predicted RT of iRT peptide GAGSSEPVTGLDAK, since it was defined to have a zero score on the iRT scale.

**Figure 3:**
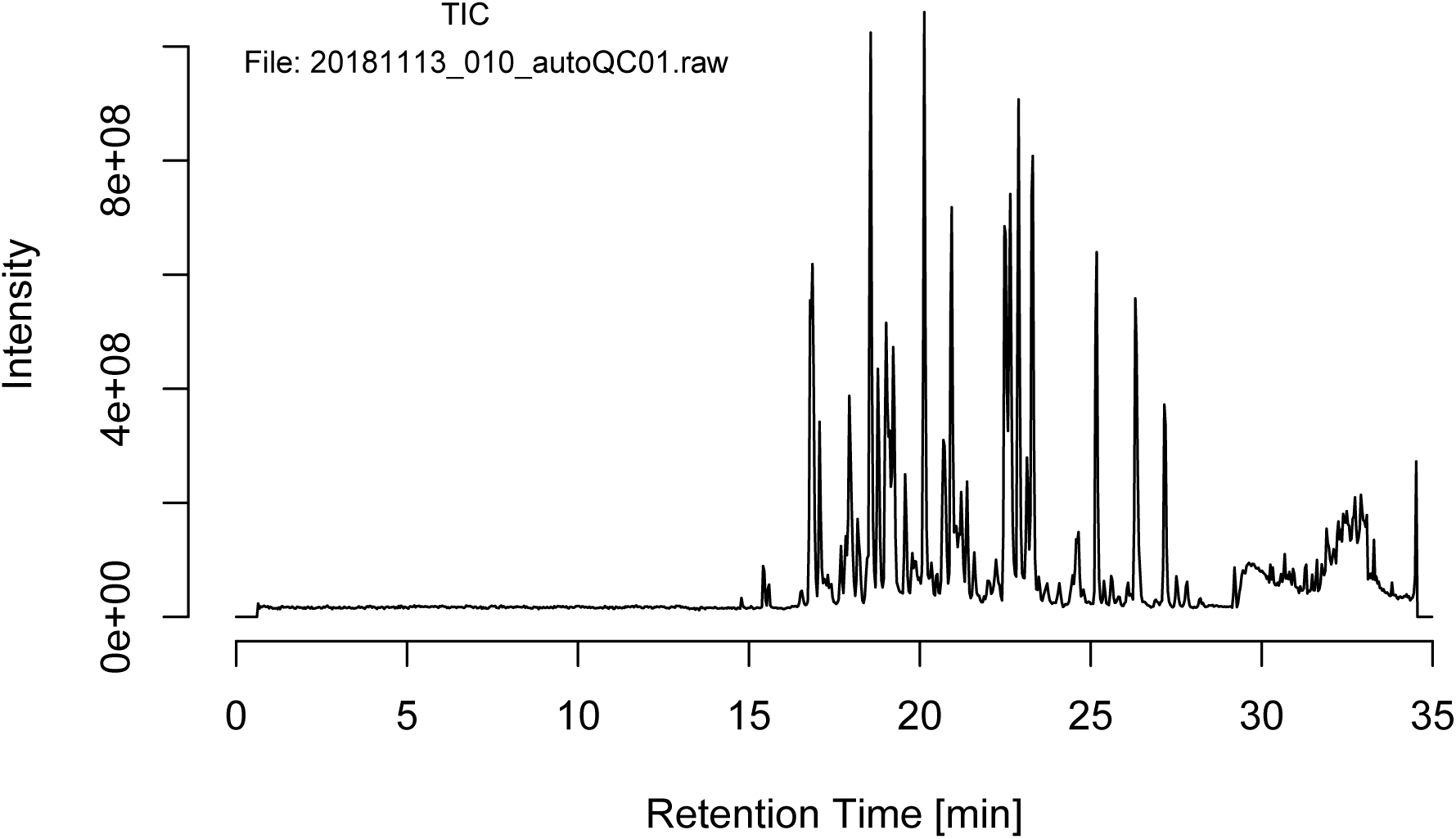
TIC.

**Figure 4:**
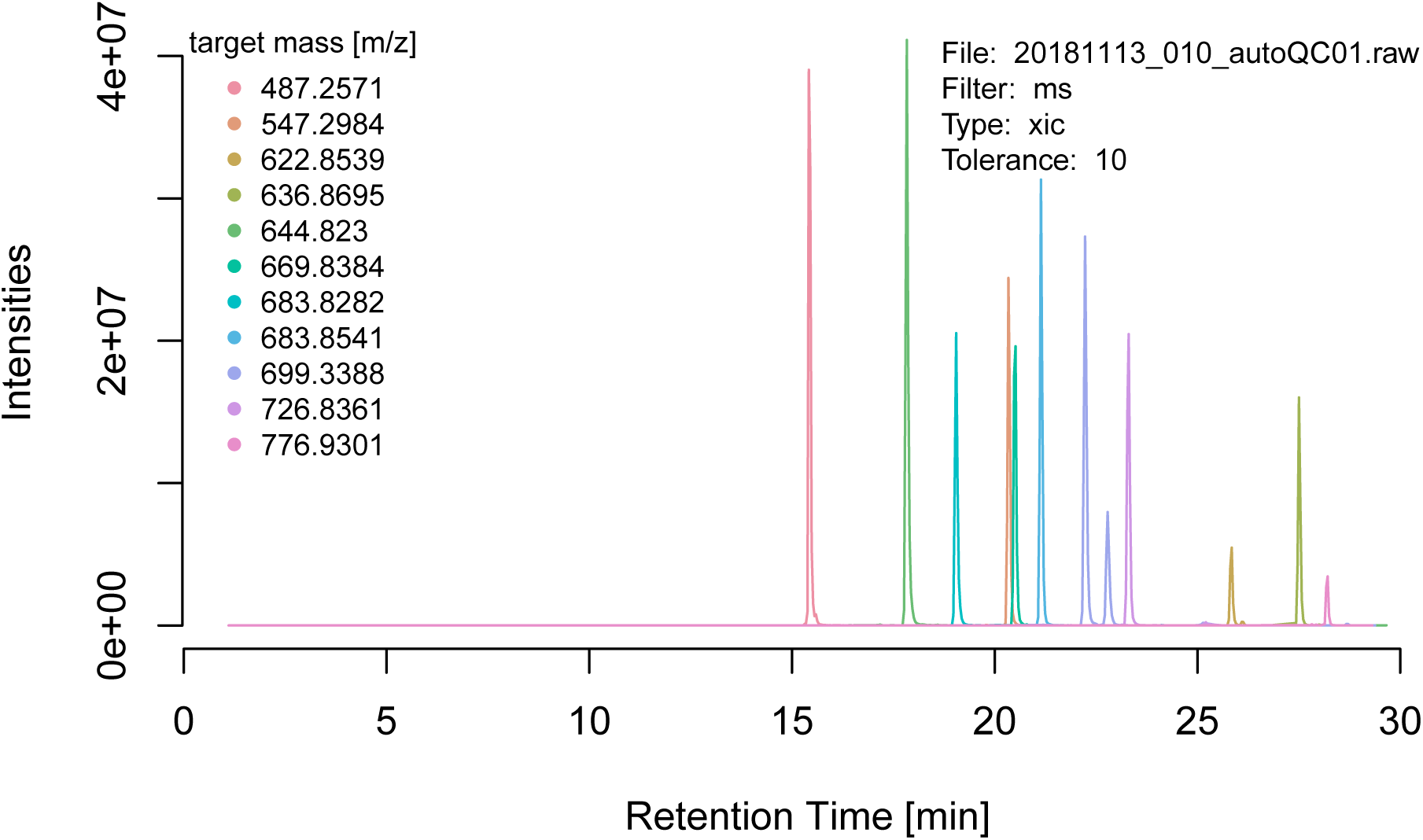
XICs for iRT peptides precursors. Each XIC was calculated using a tolerance of 10 ppm around the target mass and using only MS1 scans.

**Figure 5:**
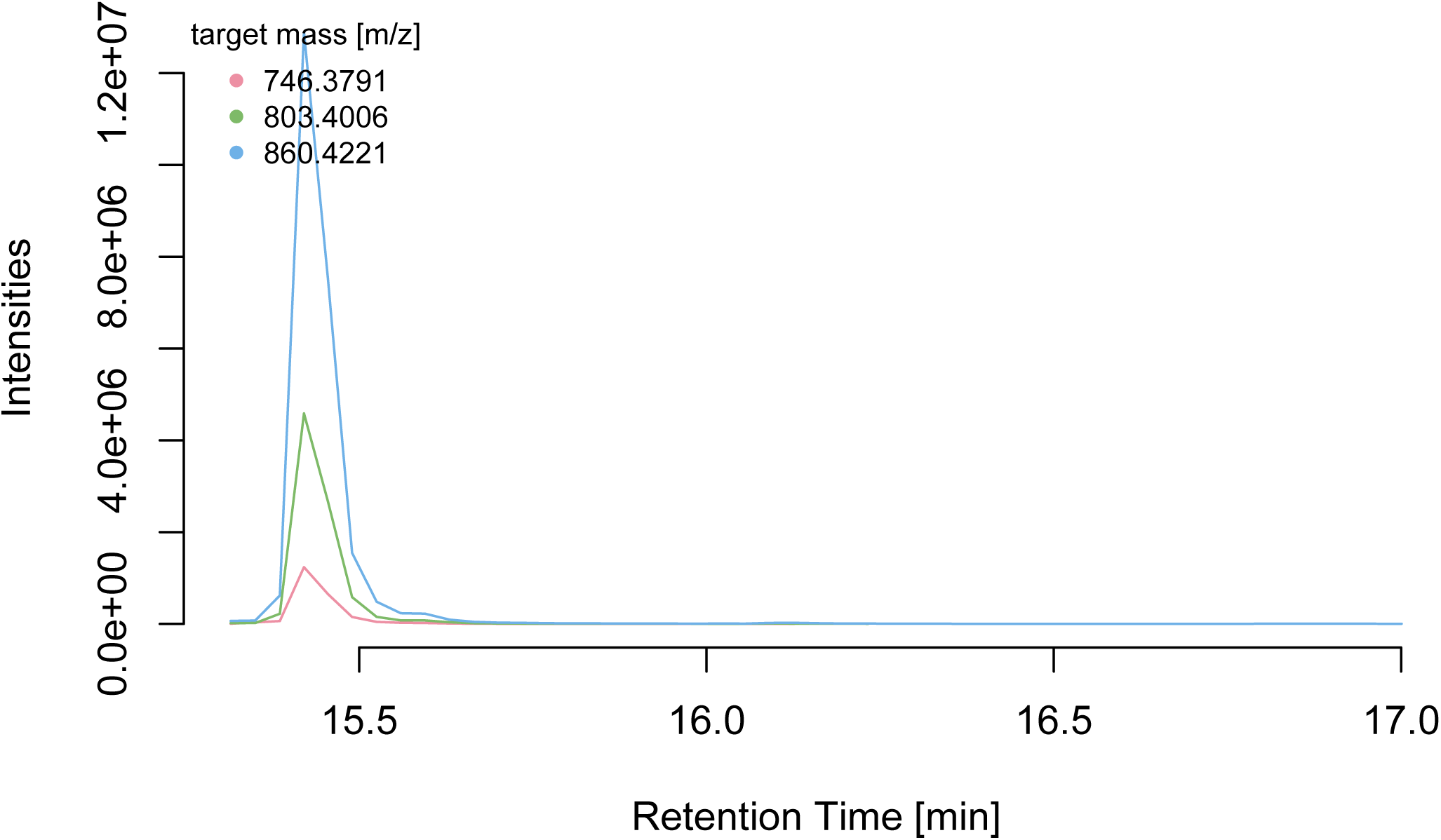
XICs for LGGNEQVTR++ fragment ions y6 to y8.

**Figure 6:**
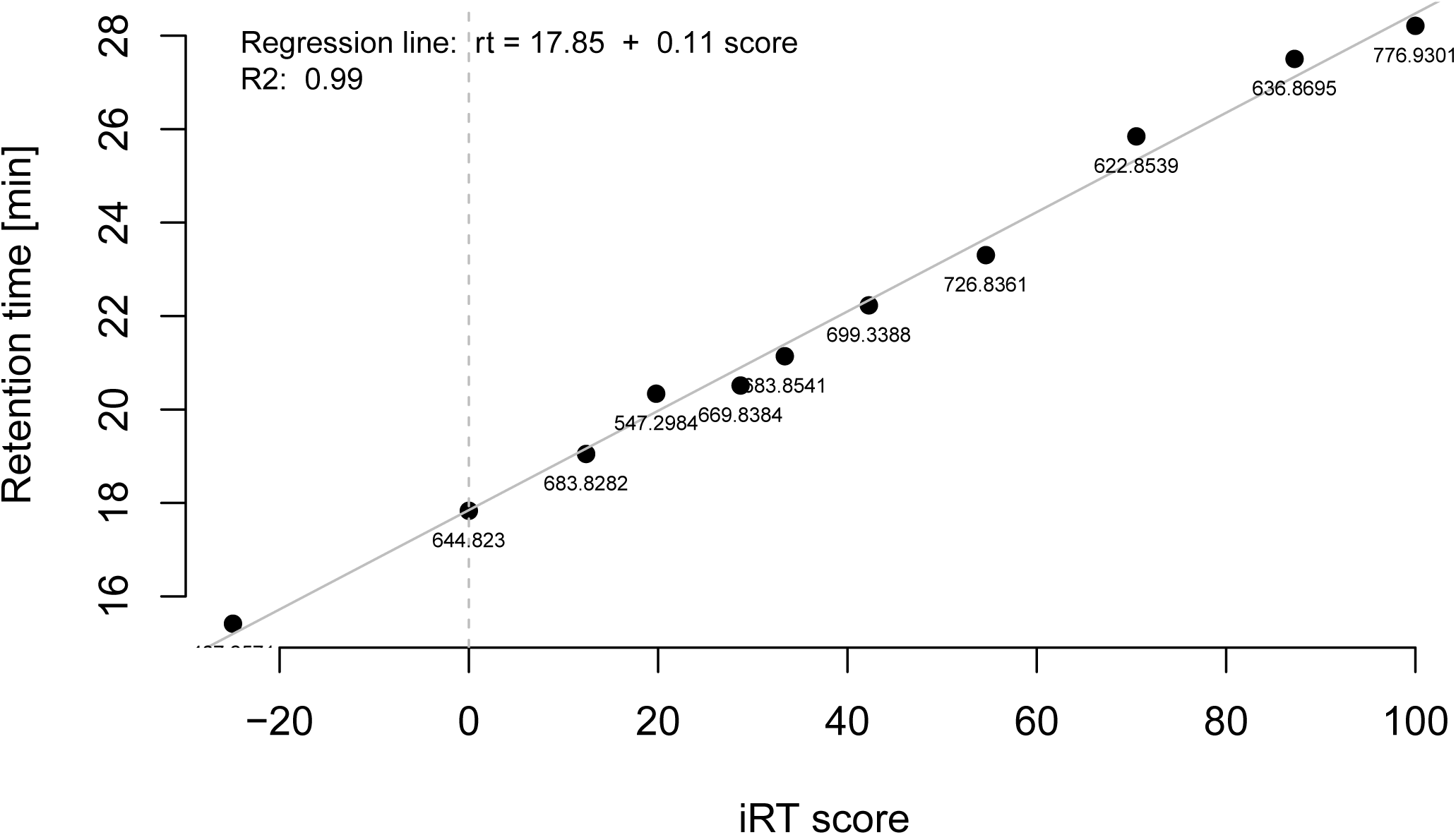
iRT score fit plot with regression line.

### Extension

An extended version of the above use cases can be found at (https://fgcz-ms.uzh.ch/~cpanse/rawR/test/functional_test.html). The web page displays spectra and iRT regression models obtained over a set of rawfiles recorded approx. every 12 hours on different orbitrap mass spectrometers at the FGCZ (some systems are not on duty anymore). The original purpose of these injections is automated longitudinal system suitability monitoring and quality control. In this case, we use the resulting rawfiles to test `rawR` functionality over different orbitrap instrument. In order to find the high-scoring MS2 scan for LGGNEQVTR++ per file, we now use a simple scoring function, implemented directly in `R` (it actually counts the number of matching y-ions), instead of running an external search engine. The web page automatically updates every 30 min using the most recent two files per system as input data. Be aware that the source code is executed in a full parallel fashion (each core processes one rawfile) on a Linux server! This shows, how scalable analysis pipelines can be constructed starting from basic building blocks (code chunks). It demonstrates that (i) `rawR`s data access mechanism works for all types of instrument models and (ii) over network attached storage.

## Conclusions

Our R package `rawR` provides direct access to spectral data stored in vendor-specific binary raw files, thereby eliminating the need for unfavorable conversion to exchange formats. Within the `R` environment, spectral data is presented by using only two non-standard objects representing data items well known to analytical scientists (spectrum & chromatogram). This design choice makes data handling relatively easy and intuitive and requires little knowledge about internal/technical details of the implementation. By using vendor API methods whenever possible, we nevertheless made sure that ease-of-use doesn’t impair performance. We also emphasized that our implementation aligns well with common `R` conventions and styles. In the near future, we plan to submit `rawR` to the BioC project and align further efforts with the R for Mass Spectrometry initiative. In particular, we hope to extend `rawR` towards the concept of exchangeable backends for data access and parallel computation. These would be necessary next steps towards big computational proteomics in `R`.

## Author contributions

The manuscript was written through contributions of all authors. All authors have given approval to the final version of the manuscript. ‡These authors contributed equally.

## Acknowledgements

We thank Lilly van de Venn for designing the `rawR` package logo. We are grateful to Jim Shofstahl for providing the `RawFileReader` .NET assembly, `C#` example code, and for answering questions during the development process of `rawR`.

## Abbreveations

MS: mass spectromerty
TIC: total ion chromatogram
XIC: extracted ion chromatogram
(FTMS): fourier-transformed mass spectrum
NSI: nano spray ionisation

## Supplements

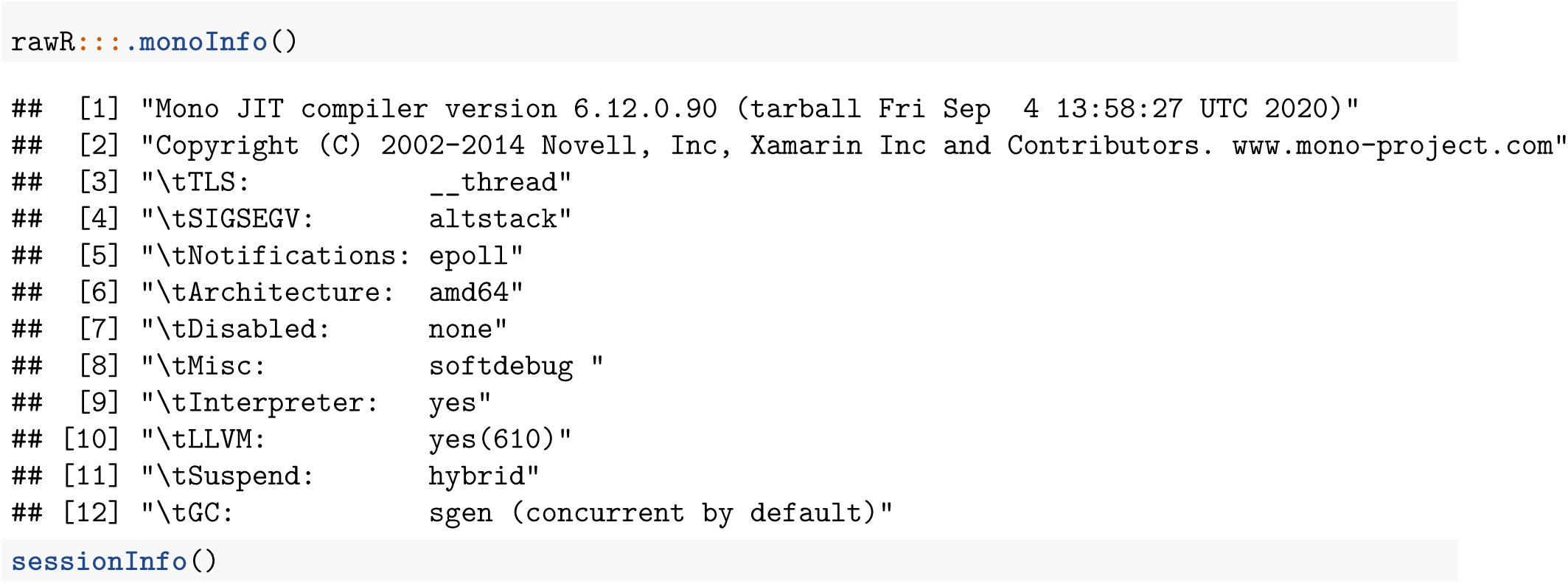

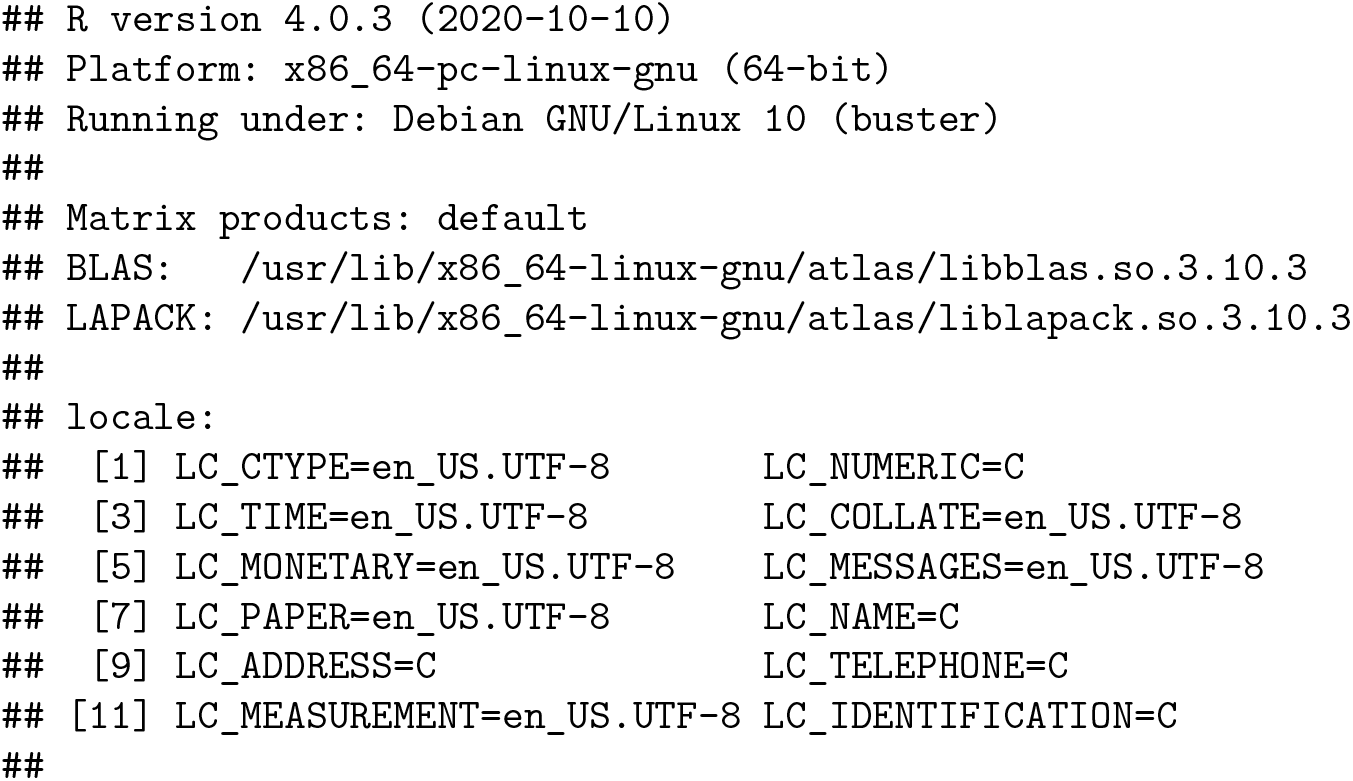

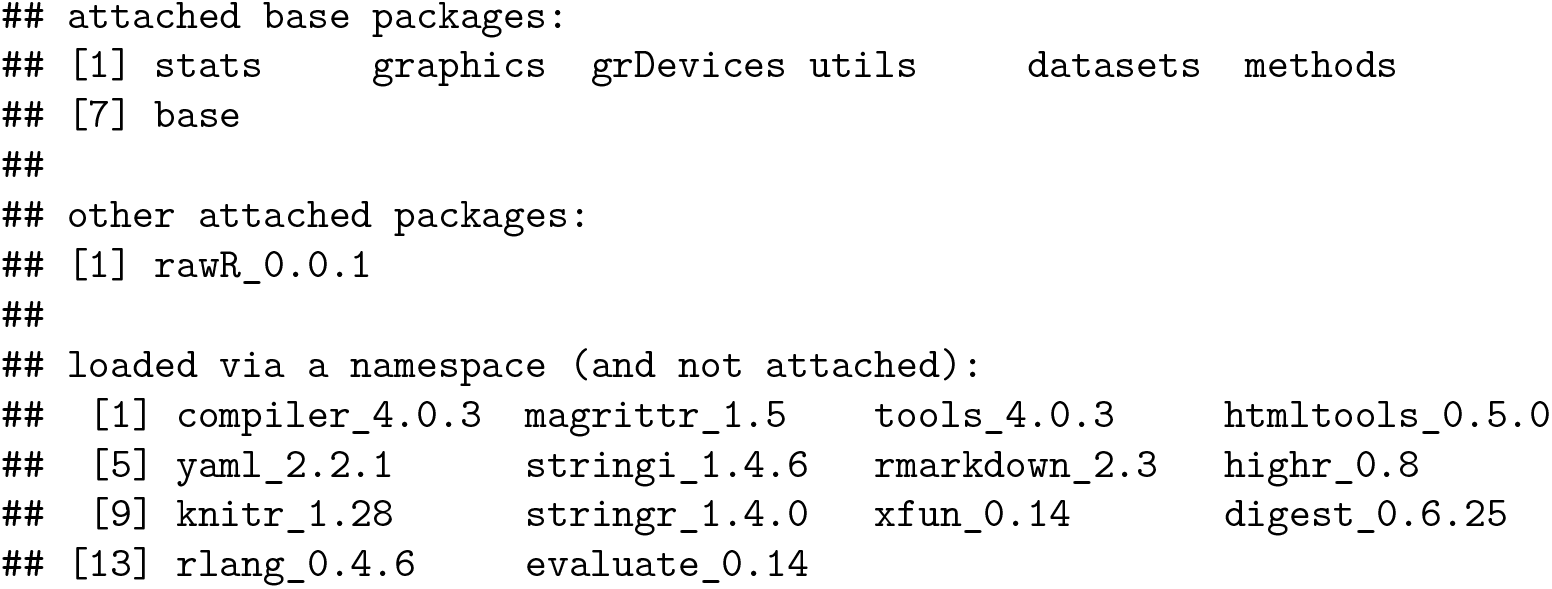

